# Beyond Metabolites: A Wearable Differential Biointerface Integrating Antibody and Aptamer Probes for the Real-Time Tracking of Proteins In Vivo

**DOI:** 10.64898/2026.03.27.714878

**Authors:** Hanjia Zheng, Fatima Shafique, Alexander S Qian, Mayank Garg, Florian Gessler, Jonathan L’Heureux Hache, Bernardo L Trigatti, Mahla Poudineh, Leyla Soleymani

## Abstract

Continuous monitoring of protein biomarkers could transform the management of acute and chronic diseases. Despite tremendous potential, wearable health monitors have remained largely limited to metabolites and small molecules. A key challenge is the limited availability of biointerfaces that reversibly track low-abundance proteins *in vivo* without user intervention. Here, we present the Differential Aptalyzer, a minimally invasive hydrogel microneedle platform for continuous monitoring of proteins in skin interstitial fluid. The platform combines high-affinity antibodies for selective target capture with aptamers for reversible electrochemical signal transduction. When integrated into a differential electrochemical chip and pulse-assisted sensor regeneration, this approach enables continuous monitoring of proteins in a wearable format. Using cardiac troponin I (cTnI) as a clinically-relevant model analyte, Differential Aptalyzer offers a broad dynamic range (0.003–0.640 ng/mL) and strong specificity against interfering proteins. Importantly, this platform reliably tracks both rising and falling exogenous cTnI levels injected into healthy mice, as well as endogenously elevated cTnI in a double-knockout mouse model of coronary artery disease, demonstrating its capability in continuous protein monitoring and identifying coronary artery disease cohorts.

## Introduction

Technologies for continuous monitoring of biomarkers are critically needed to improve patient outcomes across a range of diseases, including diabetes^1^, cardiovascular disease^2^, and cancer^3^, as well as for managing women’s health^4^ and perioperative care^5–7^. To enable the widespread adoption of these platforms in the realm of personalized medicine, they should be deployable to different settings both inside and outside the hospital, including ambulatory care, emergency departments, intensive care, and postoperative care. As such, there is a need for wearable bioanalytical tools that enable continuous biomarker profiling in a minimally-invasive manner without the need for user intervention at the point-of-need (**Fig. 1a**). Despite the need for such tools, the continuous glucose monitor is the only widely deployed and commercially available wearable technology that provides real-time and longitudinal bioanalytical data. These devices rely on transdermal electrodes functionalized with glucose oxidase to measure glucose in interstitial fluid (ISF) and have transformed diabetes management by enabling closed-loop and patient-directed care. However, this technology is fundamentally constrained by its reliance on enzymatic reactions, which restrict continuous monitoring to a narrow set of analytes such as small molecules and metabolites. As such, there remains a major technological gap for detecting other classes of biomarkers, particularly proteins that are central to addressing the needs of personalized medicine.

**Figure 1:**
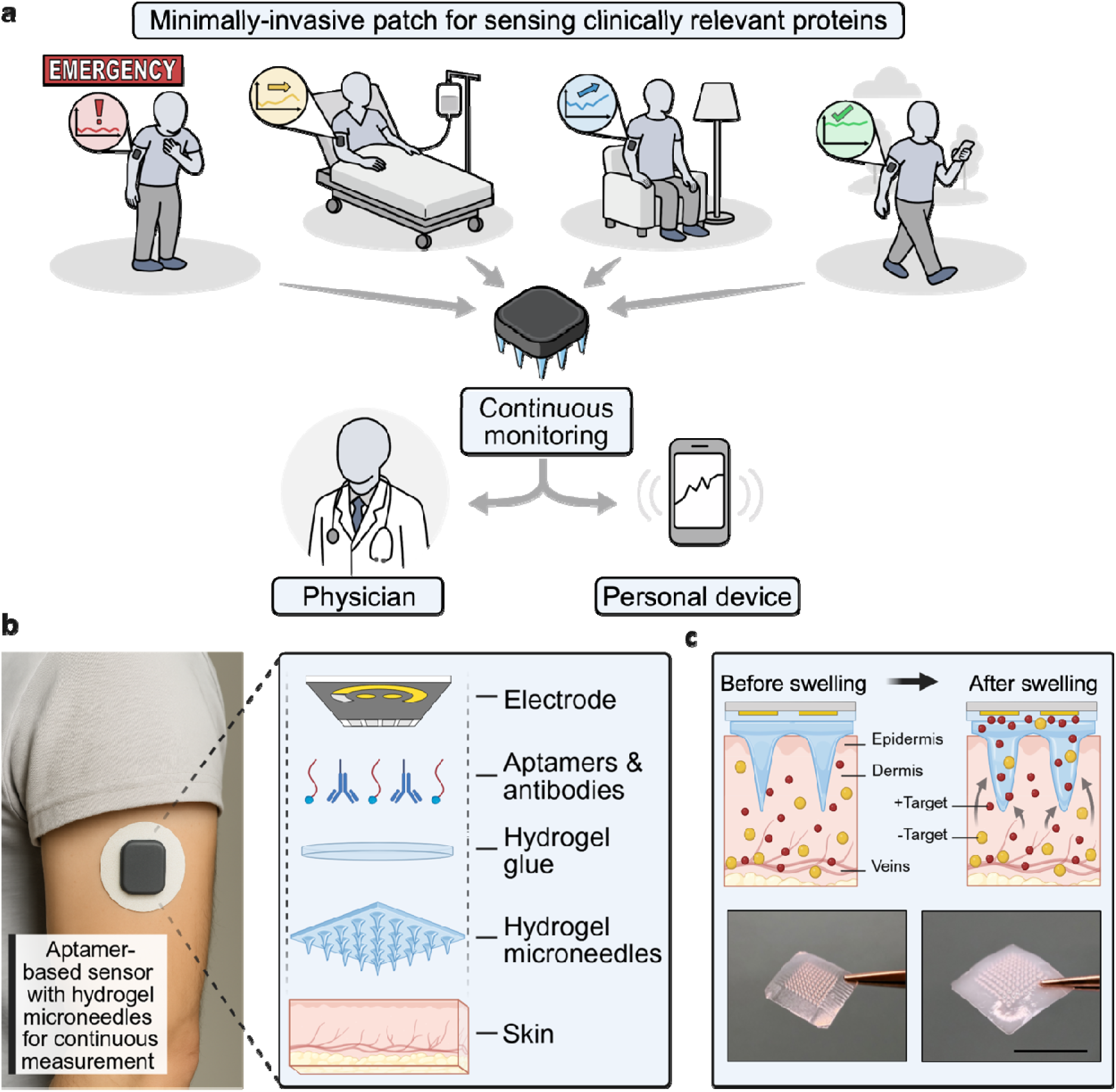
Overview of Differential Aptalyzer and its applications. **(a)** Wearable bioanalytical sensors that enable continuous, point-of-need biomarker profiling can be deployed across ambulatory settings, emergency departments, intensive care units, and postoperative care. **(b)** Schematic illustration of the components of the Differential Aptalyzer. **(c)** Top: Cross-sectional schematic illustration of the HMN patch before and after swelling, highlighting biomarker diffusion and ISF absorption. Bottom: Optical images of HMN patches before and after swelling, demonstrating patch enlargement upon swelling. Scale bar = 7 mm.

Continuous protein sensing is uniquely challenging because many clinically-relevant proteins circulate at very low concentrations (nanomolar to picomolar)^8^, necessitating high-affinity capture probes with low dissociation constants. Despite the availability of *in vitro* assays for high-sensitivity protein analysis, their translation into wearable platforms suitable for continuous monitoring encounters three fundamental barriers. First, many of such high sensitivity assays rely on multi-step workflows involving sequential washing and the addition of exogenous reagents, which preclude true real-time operation^9^. Second, high-affinity binding interactions are often associated with slow dissociation kinetics, which determine equilibration timescales and can delay sensor response, thereby limiting the ability to track decreasing biomarker levels in real-time^10–12^. Third, specimen collection for *in vitro* assays requires repeated invasive steps such as venipuncture that is challenging for many of the abovementioned settings.

To date, wearable electrochemical technologies that aim at moving beyond enzymatic sensing have either focused on using antibodies or aptamers for target capture and identification. Aptamers are nucleic acid analogues of antibodies that can be selected *in vitro* for binding a wide range of analytes, including proteins, small molecules, and ions^13^. Aptamers can be directly modified with redox reagents and engineered to undergo a reversible conformational change upon target capture, enabling both reagentless and reversible analysis^14^ to address the limitations of antibody-based systems. Nevertheless, aptamer-based biosensors reported to date for *in vivo* monitoring have largely focused on small-molecule targets in ISF, including metabolites such as glucose^15^, lactate^15^, and phenylalanine^16^, as well as pharmaceutical compounds such as tobramycin^17,18^, vancomycin^17,19^, irinotecan^20^. Recent work has demonstrated continuous *in vivo* monitoring of protein biomarkers, including inflammatory cytokines such as IL-6 and TNF-α, using aptamer-based electrochemical sensor^21^. Despite these advances, extending aptamer-based assays to *in vivo* monitoring of proteins remains challenging due to the difficulty and the time-intensive process of selecting aptamers that can both bind protein targets and undergo the necessary conformational change for signal readout^22–24^. While such conformation-changing aptamers have been developed for small molecules^15–20^, relatively few aptamers for protein targets have been translated into signaling platforms^25,26^, with representative examples including thrombin^27–29^, cardiac troponin^30^, C-reactive protein^31^, and viral proteins such as RSV-G25^32^. A limited number of antibody-based systems have also been reported for *in vivo* detection of target analytes such as cortisol^33^, pro-inflammatory cytokines^34–36^, cardiac troponins^37^, host antibodies^38^, as well as broader protein biomarkers in interstitial fluid^39^; however, these systems are not capable of truly real-time, continuous monitoring in ISF. Instead, these systems are either implemented as single-use or quasi-continuous sensors^33–36,38,39^ or have not been demonstrated to regenerate and respond to decreasing target concentrations^36,37^. This limitation is rooted in the inability of antibodies to directly generate an electrochemical signal in response to increasing and decreasing target concentration, requiring *in vitro* steps such as reagent addition and washing. Recent work on reagentless molecular pendulum sensors has begun to address the scientific barrier in protein sensing by combining double stranded DNA scaffolds that are redox labelled with biorecognition elements such as antibodies^23,40^ or aptamers^21,23,40^. By applying time-varying electric fields to drive rapid oscillation of the sensing construct, these systems can temporally resolve binding events and promote dissociation when analyte concentrations decline, thereby enabling continuous protein measurement using electrochemical readout. Nevertheless, it is not clear whether such oscillation-based signal transduction can be applied to real-life settings that involve highly active and moving human subjects.

To address the limitations of continuous protein monitoring—and building on our previously reported aptamer-based wearable platform for glucose and lactate sensing^15^—we present the Differential Aptalyzer: a minimally invasive wearable platform engineered for continuous tracking of protein biomarkers. The system overcomes the limitations of antibody or aptamer-based sensors for real-time monitoring by integrating these two biorecognition elements and harvesting their synergistic interactions. More specifically, high-affinity antibodies are used for selective protein capture, while aptamers enable reversible and reagentless electrochemical signal transduction. Importantly, the antibody/aptamer interface is linked in operation. The capture of the protein target by the antibody hinders the access of the aptamer to its small molecule target that is native in the human body. Such interactions induce a differential response that is correlated with the protein concentration.

To support wearable operation, antibody- and aptamer-functionalized electrodes were embedded within a hydrogel microneedle (HMN) patch (**Fig. 1b**). The HMNs are applied in the dry state to facilitate skin insertion and subsequently swell upon ISF uptake, forming a hydrated, porous network that enables analyte diffusion to the sensing interface while maintaining skin compatibility and user comfort (**Fig. 1b**). As a model protein, we demonstrated the Differential Aptalyzer’s capability for continuous monitoring of cardiac troponin I (cTnI) whose frequent, accessible, and rapid tracking is critical in emergency settings, intensive care, and postoperative care for the diagnosis of myocardial infarction. For example, in managing acute coronary syndromes, time-to-diagnosis directly influences revascularization strategies, emergency department crowding, and downstream resource utilization. Although high-sensitivity cTnI assays have markedly improved diagnostic accuracy, their deployment remains largely centralized, dependent on laboratory infrastructure and serial sampling protocols. Consequently, the clinical bottleneck has shifted from analytical sensitivity to turnaround time and workflow integration^41^.

To address the unmet need for frequent monitoring of cTnI with a rapid turnaround time and de-centralized workflows, we developed and comprehensively validated the Differential Aptalyzer through *in vitro* characterization, followed by *in vivo* testing in a mouse model of coronary artery disease to detect endogenously elevated cTnI. The sensor reliably distinguished animal cohorts with elevated cTnI from those with baseline levels. Furthermore, by employing pulse-assisted sensor regeneration, we monitored the ability of the sensor in tracking both increasing and decreasing concentrations of cTnI through injection studies that controllably change exogenous cTnI concentrations.

## Results and Discussion

### Differential Aptalyzer sensing mechanism

We developed the Differential Aptalyzer to address the unmet need of continuous protein tracking using a wearable platform. This assay is based on a unique biointerface that includes antibodies for the selective capture of protein targets, in this case cTnI, and aptamers for continuous signal transduction (**Fig. 2a**). Importantly, antibodies and aptamers are interlinked in operation: target capture by the antibodies and the formation of antibody/target complexes reduce the access of the aptamers to their native target, in this case, lactate. This in turn modulates the electrochemical signal, measured using square wave voltammetry (SWV), that is generated by conformational changes of the aptamers, allowing antibodies to serve as capture probes and aptamers to function as reporting probes.

**Figure 2:**
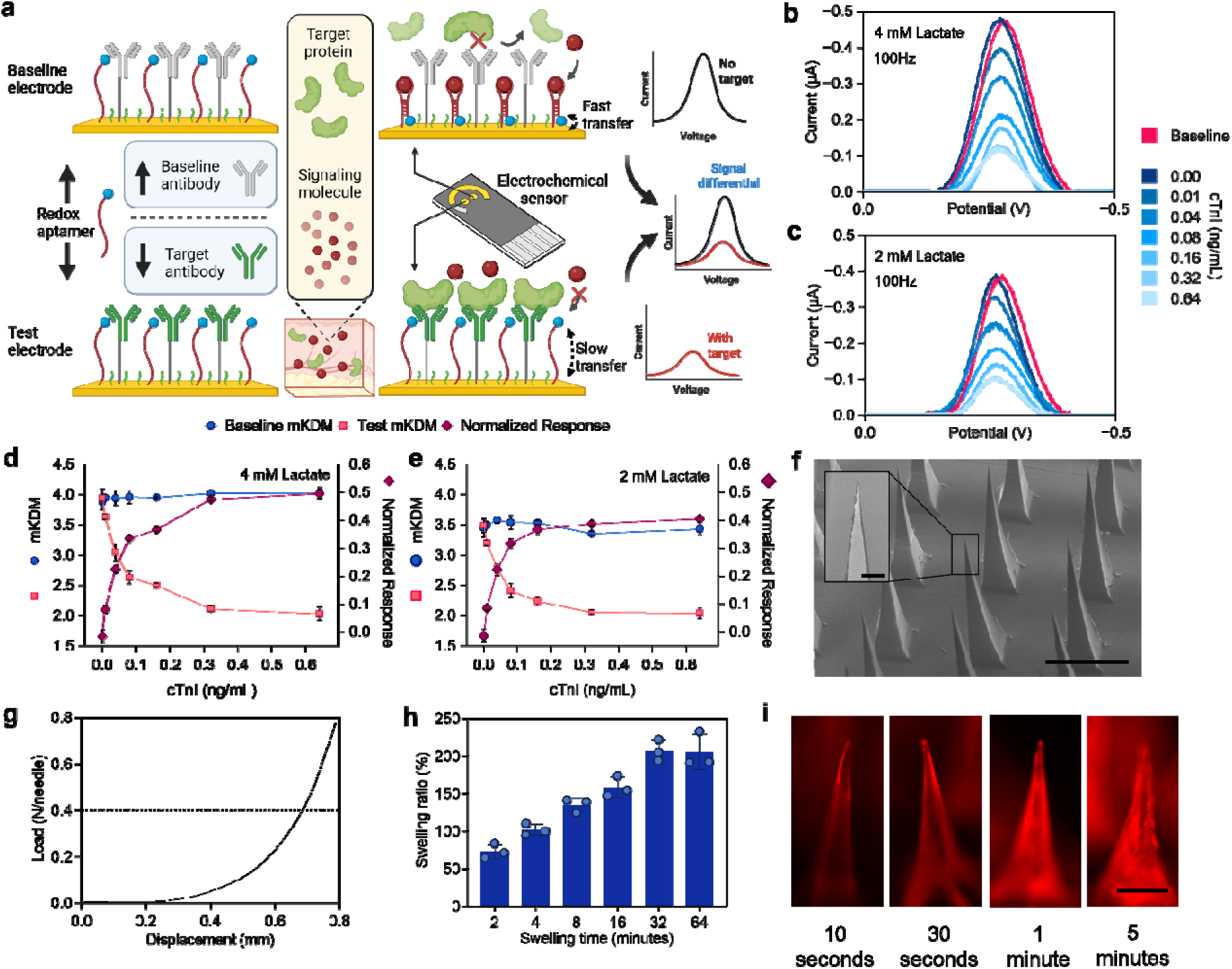
Differential Aptalyzer validation and HMN characterization. **(a)** Schematic illustration of Differential Aptalyzer signal transduction. The test electrode (anti–target antibody + redox aptamer for a signaling molecule) shows a reduced SWV peak current when target protein binding hinders access of the signaling molecule to the aptamer, suppressing the aptamer conformational change and electron transfer. In contrast, the baseline electrode (non-target antibody + the same aptamer) does not capture the target protein, enabling normal signaling-molecule binding, higher charge transfer rate, and a higher SWV peak current. The target protein concentration is quantified from the differential (test minus baseline) signal. SWV curves recorded at the test and baseline electrodes at the on-frequency (100 Hz) across varying cTnI concentrations, measured in **(b)** 4 mM and **(c)** 2 mM lactate. SWVs were recorded at the on-frequency (100 Hz) and off-frequency (25 Hz) in artificial ISF spiked with varying cTnI concentrations, measured in **(d)** 4 mM and **(e)** 2 mM lactate. All SWV measurements were performed using an on-chip Ag/AgCl reference electrode with an on-chip Au counter electrode over potential range of 0.0 to −0.5 V at 0.1 V/s and amplitude of 25 mV. The Normalized Response was calculated as. Analysis was performed on three independent chips; data are presented as mean ± standard deviation (error bars). **(f)** Scanning electron microscopy images of a HMN patch. Scale bar = 500 µm. The inset shows the magnified view of the tip of a single needle within the HMN patch. Scale bar = 20 µm. **(g)** Force-displacement curve obtained from mechanical testing of an HMN patch (n = 1). **(h)** Swelling ratio of HMN patches after application to porcine skin for different durations. Each condition was tested using three HMN patches. Data are presented as mean ± standard deviation (error bars). **(i)** Fluorescence images of a single needle in HMN arrays after application to agarose hydrogel containing 7 µg/mL Rhodamine B for different time durations. Scale bar = 150 µm. Each condition was tested using one HMN patch.

To directly correlate the concentration of the cTnI to the signals measured by the lactate aptamers, we developed a differential electrochemical chip with a test and a baseline electrode. The test electrode combines anti-cTnI antibodies with lactate aptamers; whereas, the baseline electrode combines anti-bovine serum albumin (anti-BSA) antibodies – serving as a non-specific control with no reactivity to human or rodent proteins – and lactate aptamers. The two electrodes are monitored simultaneously with the baseline electrode tracking the native lactate level and the test electrode measuring the lactate signals that are modulated by the capture of cTnI targets. Similar to differential measurement approaches used in electronic circuits^42^, the difference in signal between the test and baseline electrode is used to track the cTnI concentration (**Fig. 2a**).

The immobilization of the lactate aptamer and antibodies on the gold electrodes was confirmed using electrochemical impedance spectroscopy (EIS) and cyclic voltammetry (CV) (**Fig. S1**). We next optimized the lactate readout on the test electrode by evaluating the SWV current response across a range of frequencies, identifying 100 Hz as the on-frequency and 25 Hz as the off-frequency (**Fig. S2**). Using these parameters, SWV was used to monitor the test and baseline electrodes at two lactate concentrations (2 mM and 4 mM) (**Fig. 2b-c** and **Fig. S3**). The modified kinetic differential measurement (mKDM) was then calculated as the normalized difference in charge-transfer kinetics between two frequencies designed to maximize the signal change (equation 2)^43^. We also defined “Normalized Response” by scaling the difference between the test and baseline signals by the baseline electrode signal, and used this metric to quantify cTnI concentration, given by equation 3 (**Fig. 2de**). As hypothesized, the baseline electrode signal remains largely unchanged across varying cTnI concentrations; whereas, the test electrode exhibits a decrease in current (**Fig. 2b-c**) and the corresponding mKDM (**Fig. 2d-e**) with increasing cTnI. This trend is consistent with reduced lactate access to the aptamer layer caused by cTnI occupancy at the electrode surface.

To translate the Differential Aptalyzer into a wearable format, the differential chips were integrated into HMN patches fabricated from methacrylated hyaluronic acid (MeHA). MeHA was synthesized by methacrylating the hydroxyl groups of hyaluronic acid following our established protocol^15^, and the degree of substitution was confirmed by ^1^H NMR (**Fig. S4**). HMNs were produced using a molding process: MeHA was mixed with a photoinitiator and crosslinker, cast into microneedle molds, dried overnight, demolded, and UV-crosslinked for 45 minutes (**Fig. S5**). Scanning electron microscopy verified well-defined, sharp needle geometries (**Fig. 2f**) suitable for effective skin insertion. Mechanical performance was evaluated by compression testing, demonstrating that individual microneedles maintained structural integrity under forces exceeding 0.4 N needle^-1^, the threshold required to penetrate the stratum corneum^44^ (**Fig. 2g**). Swelling behavior was assessed using porcine ear skin model to mimic the human skin environment following our previous protocols^15^. Patches were applied to the porcine skin, and swelling ratios were quantified by comparing the weight of patches before and after application (equation 1). The HMNs reached maximal swelling within ∼30 minutes of application (**Fig. 2h**). Additionally, fluorescence imaging showed that the swollen HMNs begin to absorb the surrounding solution within 1 minute of application (**Fig. 2i**). Accordingly, we expect rapid filling of the microneedle matrix with ISF biomarkers, within the first minute of HMN insertion.

### *In vitro* validation of the integrated cTnI Differential Aptalyzer

Having established the Differential Aptalyzer assay and HMN patches, we assembled an integrated wearable cTnI Differential Aptalyzer by coupling antibody/aptamer-functionalized electrodes to HMN patches, following our established procedure^15^ (**Fig. 3a**). To verify whether large biomolecules (e.g., proteins) can permeate the HMN network, we evaluated the ability of redox labelled-BSA deposited on the HMN patch to be detected on the electrode surface. Importantly, BSA is larger (66 kDa) than cTnI (24 kDa), making it a stringent surrogate model for assessing protein diffusion through the HMN matrix. SWV measured at the electrode surface showed a progressive increase in signal as the microneedles swelled, reaching a plateau at 30 minutes (**Fig. S6**), supporting that cTnI can diffuse through the HMNs and reach the electrode for detection.

**Figure 3:**
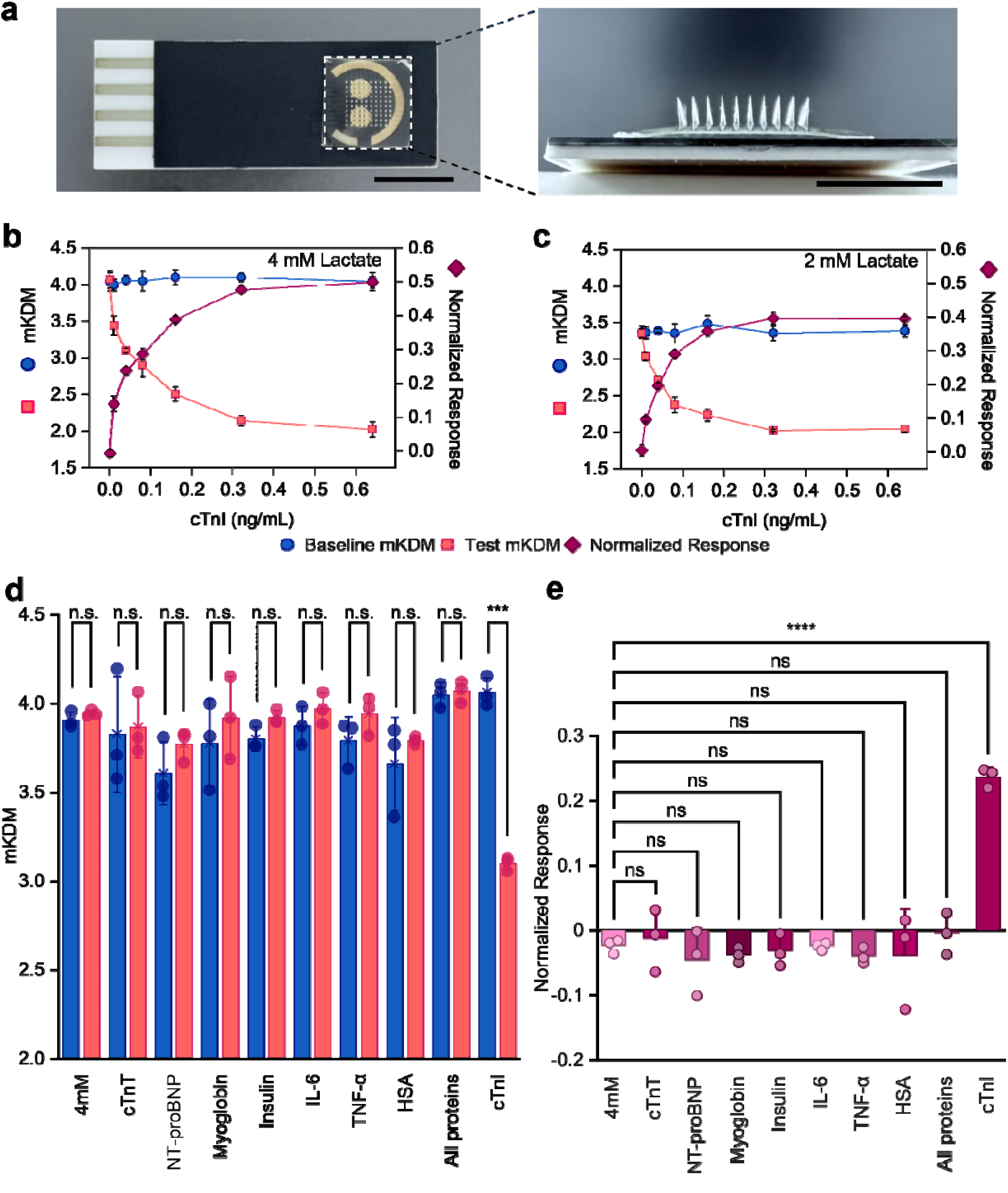
*In vitro* validation of the integrated cTnI Differential Aptalyzer. **(a)** Optical image of the fully integrated cTnI Differential Aptalyzer (left, scale bar = 1 cm), and a zoomed-in side view of the attached HMN patch (right, scale bar = 0.5 cm. The integrated device was exposed to artificial ISF spiked with varying cTnI concentrations at fixed lactate levels of 4 mM **(b)** and 2 mM **(c)**. All SWV measurements were performed using an on-chip Ag/AgCl reference electrode with an on-chip Au counter electrode over 0.0 to −0.5 V with a scan rate of 0.1 V/s and voltage amplitude of 25 mV. Measurements at 25 Hz and 100 Hz frequency were used to calculate mKDM and derive the subsequent Normalized Response. Analysis was performed on three independent chips; data are presented as mean ± standard deviation (error bars). **(d)** The specificity of the integrated cTnI Differential Aptalyzer for cTnI (0.04 ng/mL) was evaluated against common ISF interferents (0.2 ng/mL cTnT, 0.2 ng/mL NT-proBNP, 1.2 µg/mL myoglobin, 700 pM insulin, 5 ng/mL IL-6, 5 ng/mL TNF-α, 13 g/L HSA, and a mixture of all interferents) in 4 mM lactate solution. The mKDM values obtained under each interferent condition were compared with the target condition using two-way ANOVA with Bonferroni’s multiple-comparisons test. **(e)** Normalized Responses derived from data in (d). Statistical significance was assessed using ordinary one-way ANOVA with Dunnett’s multiple-comparisons test. (ns, not significant; *P < 0.05; **P < 0.01; ***P < 0.001; ****P < 0.0001).

We next quantified the cTnI Differential Aptalyzer response across cTnI concentrations of 0.00–0.64 ng/mL at fixed lactate levels of 2 and 4 mM. As shown in **Fig. 3b-c**, mKDM values are measured for both the baseline and test electrodes, and the Normalized Response is derived accordingly. The Differential Aptalyzer demonstrates a limit of detection of 0.0048 and 0.0027 ng/mL (4.8 and 2.7 ng/L) at 2 and 4 mM lactate concentrations, respectively. This limit of detection is slightly higher than that reported for a commercial point-of-care high sensitivity cTnI assay^45^ (1.05 ng/L in plasma), which is not applicable to continuous monitoring. The limit of quantification was determined to be 0.013 and 0.052 ng/mL (13 and 5.2 ng/L) at 2 and 4 mM lactate concentrations, respectively.

Specificity was then evaluated in 4 mM lactate against potential interferents, including cardiac troponin T (cTnT), NT-proBNP (two other clinically relevant cardiac biomarkers), insulin, IL-6, human serum albumin (HSA), TNF-α, and a mixture containing all interferents. mKDM measurements for both baseline and test electrodes (**Fig. 3d**), as well as the resulting Normalized Response (**Fig. 3e**), showed negligible signals in the presence of interferents and a strong response to cTnI, demonstrating excellent assay specificity for the cTnI Differential Aptalyzer.

### *In vivo* tracking with the cTnI Differential Aptalyzer

To enable the cTnI Differential Aptalyzer to track both rising and falling target protein concentrations, we developed a pulse-assisted sensor regeneration protocol in which an electric field is applied to promote target dissociation. This approach is particularly important for tracking decreasing target concentrations. Since antibodies bind their targets with high affinity, sensor recovery is typically slower when target levels drop than when they rise. Accordingly, after each measurement, a brief pulsed potential was applied to the electrode to disrupt antibody-target complexes and accelerate signal recovery. We evaluated a range of pulse amplitudes and durations and identified conditions that effectively regenerate the sensor response (**Fig. S7**). When no pulse was applied, the test electrode failed to fully return to its initial signal level after cTnI removal. In contrast, the baseline electrode—where cTnI capture is not expected—showed a reversible response even without pulsing. Therefore, we applied the reset pulse only to the test electrode to regenerate the anti-cTnI antibody layer and optimized the pulse parameters to 0.3 V for 15 s followed by −0.3 V for 15 s. We validated this strategy *in vitro* using a stepwise sequence of solutions: artificial ISF (buffer), 4 mM lactate, 4 mM lactate with 0.08 ng/mL cTnI, 4 mM lactate, and finally artificial ISF. Immediately before each new solution was introduced, a reset pulse was applied (**Fig. 4a**). As expected, introducing 4 mM lactate increased the signal on both the test and baseline electrodes. Upon addition of cTnI, the baseline electrode remained unchanged; whereas, the test electrode signal decreased. Importantly, upon switching back to 4 mM lactate, the baseline electrode response remained unchanged, as expected, while applying the reset pulse to the test electrode rapidly shifted its signal back toward pre-exposure levels (**Fig. 4b**). These results demonstrate that pulse-assisted sensor regeneration enables the cTnI Differential Aptalyzer to reliably track both increasing and decreasing target concentrations.

**Figure 4:**
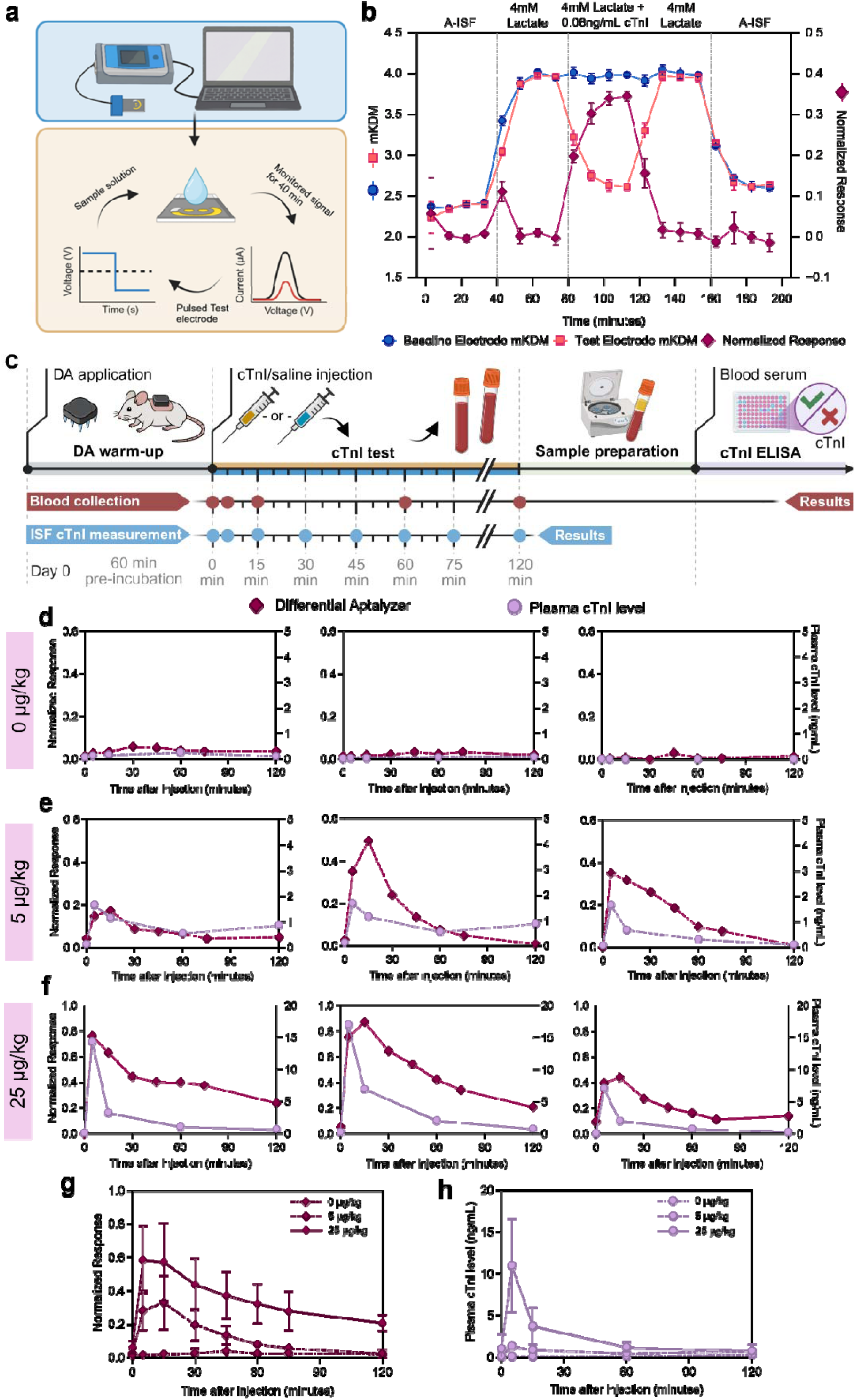
*In vivo* tracking of fluctuating cTnI concentrations. **(a)** Schematic illustration of the optimization protocols for pulsed-assisted regeneration. **(b)** In-solution validation of the pulse-assisted regeneration protocol using the cTnI Differential Aptalyzer. The device was challenged with a stepwise sequence of solutions: artificial ISF (buffer), 4 mM lactate, 4 mM lactate + 0.08 ng/mL cTnI, 4 mM lactate, and finally artificial ISF. Before each solution exchange, a pulse of 0.3 V for 15 s followed by −0.3 V for 15 s was applied to the test electrode. Analysis was performed on three independent chips; data are presented as mean ± standard deviation (error bars). **(c)** Schematic illustration of the *in vivo* study design for tracking dynamic cTnI levels. Two wearable cTnI Differential Aptalyzers were applied to the dorsal skin of mice for 60 minutes, followed by baseline cTnI measurement and baseline blood collection for plasma analysis. Mice then received cTnI injections (0, 5, or 25 µg/kg), and cTnI dynamics were monitored with the cTnI Differential Aptalyzer at 0, 5, 15, 30, 45, 60, 75 and 120 minutes post-injection, and by ELISA of plasma from heparinized blood collected from the facial vein at 0, 5, 15, 60, and 120 minutes. cTnI profiles from three mice following injections of 0 µg/kg (**d**), 5 µg/kg (**e**), and 25 µg/kg (**f**), measured by the cTnI Differential Aptalyzer and quantified in plasma using a benchtop ELISA. Summary of the normalized cTnI Differential Aptalyzer (**g**) or plasma cTnI (**h**) results across the three dosing groups (n = 3).

The *in vivo* performance of cTnI Differential Aptalyzer for tracking dynamic cTnI fluctuations was evaluated in a mouse model. To generate controlled cTnI kinetics, healthy mice were administered either saline (0 µg/kg) or cTnI at two doses (5 µg/kg and 25 µg/kg), while the cTnI Differential Aptalyzer—operated using the pulse-assisted sensor regeneration protocol—continuously monitored cTnI. Before measurements, two cTnI Differential Aptalyzers were applied to the shaved dorsal skin of mice and left in place for 60 minutes to allow the HMNs to fully swell. cTnI Differential Aptalyzer measurements were acquired at 0, 5, 15, 30, 45, 60, 75 and 120 minutes after injection. In parallel, blood samples were collected at 0, 5, 15, 60, and 120 minutes and analyzed for plasma cTnI using enzyme-linked immunosorbent assay (ELISA; **Fig. 4c**). In the saline group (**Fig. 4d**), the cTnI Differential Aptalyzer signal remained stable with no measurable change. In contrast, in mice injected with 5 µg/kg cTnI (**Fig. 4e**) and 25 µg/kg cTnI (**Fig. 4f**), the cTnI Differential Aptalyzer captured both the rise and subsequent decline in cTnI over time. These temporal profiles were consistent with cTnI concentrations measured by ELISA from serial blood draws. Notably, the cTnI Differential Aptalyzer also resolved dose-dependent differences: as expected, the 25 µg/kg injection produced a larger response than 5 µg/kg (**Fig. 4g-h**).

### *In vivo* validation of cTnI Differential Aptalyzer using a mouse model of coronary artery disease

To assess the feasibility of the cTnI Differential Aptalyzer in detecting elevated cTnI concentrations associated with myocardial infarction, we evaluated its performance in a mouse model of coronary artery disease. We employed a genetically modified knockout line of mice, which lacks expression of both low-density lipoprotein (LDL) receptor (LDLR) and the high-density lipoprotein (HDL) receptor, scavenger receptor class B type 1 (SR-B1). On a standard low fat, low cholesterol diet, these double knockout (DKO) mice do not develop atherosclerosis and do not exhibit significant cardiovascular pathology^46,47^. However, when they are fed a high-fat/high-cholesterol, cholate-containing (HFHC) diet (15% fat, 1.25% cholesterol, 0.5% cholate), they rapidly (within 3 weeks) develop atherosclerotic plaques in coronary arteries, with a significant proportion of coronary artery cross sections showing complete occlusion with atherosclerotic plaques (**Fig. 5a** and **Fig. S8**^46^). These HFHC diet-fed DKO mice show signs of myocardial infarction, indicated by greater myocardial damage and fibrotic scarring in Masson Trichrome stained transverse sections of the myocardium compared to HFHC diet-fed SR-B1 heterozygous control mice (**Fig. 5b**). These phenotypes are consistent with other reports of spontaneous or atherogenic diet induced occlusive coronary artery atherosclerosis, myocardial fibrosis and cardiac dysfunction in SR-B1 knockout or mutant strains of mice on atherogenic apolipoprotein E (ApoE) or LDLR mutant backgrounds ^32,34–39^. SR-B1−/− LDLR−/− DKO mice fed the HFHC diet for 19 days of time have previously been reported to exhibit elevated levels of plasma cTnI, consistent with myocardial damage associated with myocardial infarction^47^. We tested two groups: HFHC-fed control mice (SR-B1+/− LDLR−/−, n = 8) and HFHC-fed DKO mice (SR-B1−/− LDLR−/−, n = 14) (**Fig. 5c**). At the end of the 20-day diet period, mice were prepared for cTnI measurement. Under anesthesia, two cTnI Differential Aptalyzers were applied to the shaved dorsal skin, and signals were recorded after a 60-minute stabilization period to allow HMN swelling (**Fig. 5d-e**). Measurements were then collected over 60 minutes, and the average signal was plotted. Notably, because this study aimed to assess endpoint cTnI levels rather than track short-term fluctuations, the pulse-assisted regeneration protocol was not used and an average signal was recorded. After acquiring the cTnI Differential Aptalyzer measurement, mice were then euthanized and blood was collected for cross-validation by conventional cTnI ELISA. Successful microneedle insertion was confirmed by the visible microneedle imprints on the skin after device removal **(Fig. S9)**. This is in line with our previous studies using H&E staining that verified reliable skin penetration by MeHA-based HMNs^15^.

**Figure 5:**
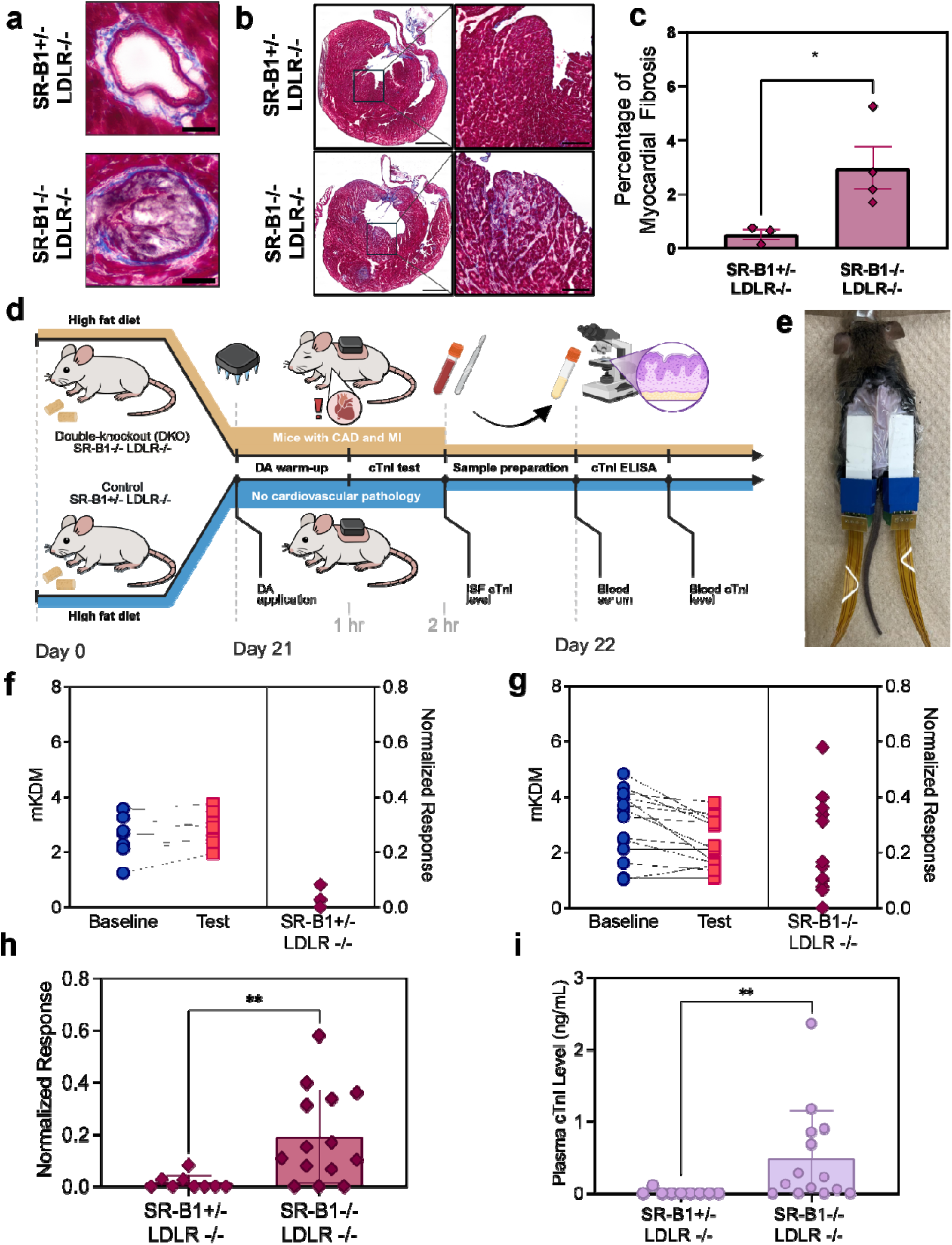
*In vivo* validation of cTnI Differential Aptalyzer using a mouse model of coronary artery disease. **(a)** Representative images of Masson’s Trichrome stained coronary arteries from SR-B1+/−LDLR−/− and SR-B1−/−LDLR−/− mice. **(b)** Representative images of transverse cryosections (scale bar, 1 mm) and corresponding close-ups of myocardium (scale bar = 200 µm) from SR-B1+/−LDLR−/− and SR-B1−/−LDLR−/− mice. Red colored staining indicates healthy myocardium, whereas blue/purple colored staining indicates the presence of collagen. **(c)** Quantification of myocardial fibrosis expressed as the percentage of blue-stained area relative to the total myocardium cross-sectional area, measured from one cross-section per mouse approximately 900 µm from the aortic annulus (n = 3−4). **(d)** Schematic illustration of the protocol used to establish the DKO mouse (SR-B1−/−LDLR−/−) and the control mouse (SR-B1+/− LDLR−/−) model, and the workflow for *in vivo* validation of the cTnI Differential Aptalyzer. **(e)** Photograph of a mouse with two cTnI Differential Aptalyzers applied to its dorsal skin for measurement. mKDM responses of individual baseline and test electrodes after application in control (**f**, n = 8) and DKO mice (**g**, n = 14). Baseline and test data points from the same device are connected using a solid line; Normalized Responses from test and baseline electrodes for each chip are shown on the right axis. Two cTnI Differential Aptalyzers were applied to each mouse for measurement (n = 2). **(h)** Normalized Response of the cTnI Differential Aptalyzer in DKO and control mice. **(i)** Plasma cTnI levels measured by benchtop ELISA in control (n = 8) and DKO mice (n = 14). Data are presented as mean and the error bars show standard deviation. Statistical significance was assessed using Mann-Whitney U test (ns, not significant; *P < 0.05; **P < 0.01; ***P < 0.001).

In control mice, where no cTnI elevation was anticipated, baseline and test electrodes yielded indistinguishable mKDM responses with a normalized response near 0 (**Fig. 5f**). By contrast, in HFHC-fed DKO mice, the test electrode exhibited a decreased signal compared to the baseline electrode, yielding a normalized response > 0.2 (**Fig. 5g**). Importantly, cTnI Differential Aptalyzer responses between baseline and test electrodes are consistent with ELISA measurements from blood samples, which confirm elevated cTnI levels in the HFHC diet fed SR-B1−/−LDLR−/− group and undetectable levels in the SR-B1+/−LDLR−/− control group (**Fig. 5h-i**). Notably, we only observe elevated cTnI in about half of the SR-B1−/−LDLR−/− mice, both in ELISA and cTnI Differential Aptalyzer results, but we see cardiac damage in all of them. This may reflect the stochastic nature of the onset of myocardial infarction resulting from occlusive coronary artery atherosclerosis, and the expected transient nature of cTnI release into plasma and ISF versus the permanent nature of the myocardial damage^48^. Together, these results demonstrate the feasibility of *in vivo* cTnI detection and its concordance with standard assays in a preclinical model of coronary artery disease.

## Conclusion

We created the Differential Aptalyzer, a minimally invasive wearable platform designed to address a long-lasting challenge in next-generation bioanalytical wearables: continuous, reagentless monitoring of low-abundance protein biomarkers in ISF. By integrating high-affinity antibodies for selective target capture with aptamers for reversible electrochemical transduction, the Differential Aptalyzer overcomes key limitations of existing antibody- or aptamer-only approaches.

We validated the platform through comprehensive *in vitro* characterization and translated it into a wearable platform by its integration with MeHA-based HMN patches. These patches provide a skin-compatible, hydrated transport matrix for biomarker diffusion to the sensing interface. The integrated device, cTnI Differential Aptalyzer, demonstrates selective detection of cTnI against clinically relevant interferents and achieves clinically-relevant sensitivity under fixed lactate backgrounds. To enable true longitudinal tracking, we established a pulse-assisted sensor regeneration protocol that accelerates the dissociation of antibody-bound targets and restores sensor responsiveness, allowing it to track both increasing and decreasing protein concentrations—an essential requirement for dynamic biomarker monitoring.

Importantly, cTnI Differential Aptalyzer performs robustly *in vivo*. In controlled injection studies, the wearable sensor captures dose-dependent rises and subsequent declines in cTnI, with temporal trends consistent with ELISA measurements from serial blood sampling. Beyond exogenous modulation, the platform also distinguishes animals with endogenously elevated cTnI in a mouse model of coronary artery disease, demonstrating concordance with gold-standard assays in a physiologically relevant setting. These results establish the Differential Aptalyzer as a practical strategy for continuous monitoring of a low abundance protein in ISF and a promising foundation for wearable profiling of clinically actionable protein biomarkers.

Beyond analytical validation, the clinical significance of this platform lies in its potential to shift how cardiac biomarkers are deployed. High-sensitivity cTnI assays have largely resolved the challenge of analytical detection^49^. However, in many healthcare systems, the remaining bottleneck is diagnostic latency driven by centralized laboratory workflows, serial sampling and processing delays^50^. A wearable platform capable of continuous cTnI monitoring introduces a fundamentally different paradigm: rather than episodic laboratory confirmation, myocardial injury could be assessed dynamically and near-continuously. Such monitoring could shorten the interval between clinically relevant concentration changes and actionable decisions, potentially reducing emergency department dwell time or enabling earlier escalation to invasive management.

Future work on the cTnI Differential Aptalyzer will include real time measurements in DKO versus control mice to determine the onset for myocardial infarction and to evaluate its suitability for early detection. We will also focus on expanding the approach to additional protein targets, including those related to myocardial infarction and heart failure, by pairing appropriate capture antibodies with redox aptamer reporters.

## Material & Methods

### Materials

Methacrylate anhydride (MA), acrylamide (MBA), Irgacure 2959 (2-hydroxy-4′-(2-hydroxyethoxy)-2-methylpropiophenone, photo initiator, PI), deuterium oxide, human cTnI (648480), and cysteamine hydrochloride, dimethylformamide (DMF), human recombinant insulin and human serum albumin (HSA) were purchased from Sigma-Aldrich. Additional reagents obtained from Sigma-Aldrich included acetone, ethanol, 2-propanol, sulfuric acid, sodium chloride (NaCl), potassium chloride (KCl), magnesium chloride (MgCl), sodium hydroxide (NaOH), sodium L-lactate, (±)-sodium 3-hydroxybutyrate, sucrose, lactose, D-(+)-glucose, glycine, L-alanine, uric acid, ascorbic acid, Bovine serum albumin (BSA), tris(2-carboxyethyl)phosphine hydrochloride (TCEP), potassium hexacyanoferrate(II) trihydrate (K_4_Fe(CN)_6_·3H_2_O), MES, phosphate buffer solution (1.0 M, pH 7.4), magnesium sulfate (MgSO_4_), sodium phosphate monobasic monohydrate (NaH_2_PO_4_) and potassium ferricyanide (K_3_Fe(CN)_6_). 4-(2-hydroxyethyl)-1-piperazineethanesulfonic acid (HEPES) were purchased from BioShop. The pharma-grade sodium hyaluronic acid (HA, molecular weight of 300 kDa) was obtained from Bloomage Co., Ltd. (China). Microneedle molds were purchased from Micropoint Technology (Signs). Anti-Cardiac cTnI antibody (ab47003), recombinant human cTnI protein (ab283299), Anti-BSA antibody (AB186531), recombinant human Cardiac Troponin T (ab316587), and Colorimetric L-Lactate assay kits (ab65331) were purchased from Abcam. 1-Ethyl-3-[3-dimethylaminopropyl]carbodiimide hydrochloride (EDC), N-hydroxysuccinimide (NHS), Alexa Fluor™ 488 Microscale Protein Labeling Kit, phosphate-buffered saline (PBS, 10×, pH 7.4), and 3.5 kDa dialysis membranes were purchased from Thermo Fisher. Additional proteins used for specificity testing including recombinant human TNF-α, recombinant human IL-6, recombinant human Myoglobin, recombinant human NT-proBNP were purchased from RayBiotech. All oligonucleotides were purchased from Integrated DNA Technologies (IDT) and purified in house using standard 10% denaturing (8 M urea) polyacrylamide gel electrophoresis (dPAGE).

### MeHA synthesis

The synthesis of MeHA was adapted and modified from previously established protocols^15^. Briefly, 2 g of hyaluronic acid was dissolved in Milli-Q water and stirred overnight at 4 °C. Subsequently, 3 mL of MA was added to the solution, followed by 6.5 mL of 5 M NaOH to maintain the pH between 8 and 9 throughout the reaction. The mixture was stirred continuously at 4 °C overnight. The resulting MeHA was then precipitated using 100% acetone and washed three times with 100% ethanol. The precipitate was redissolved in 200 mL of Milli-Q water and subjected to dialysis against deionized water for five days. Finally, the dialyzed MeHA solution was frozen at −20 °C and lyophilized to obtain the purified MeHA powder.

### MN fabrication and SEM imaging

To fabricate HMNs, 66.6 mg of MeHA, 1.3 mg of MBA, and 1.3 mg of PI were dissolved in 1 mL of Milli-Q water. Next, 300 µL of the MeHA solution was dispensed onto the mold and degassed under vacuum for 5 minutes. After drying overnight at room temperature, the HMN array was demolded and crosslinked under UV light for 40 minutes. The crosslinked HMNs were then trimmed and prepared for subsequent measurements. The crosslinked HMN patches were examined using scanning electron microscopy to evaluate the needle quality and morphology. Before imaging, the HMNs were coated with a 2 nm layer of gold. SEM imaging was then performed using HITACHI SU8700 field emission scanning electron microscope. The microneedles of the fabricated HMN patch measured 850 μm in height, 250 μm in base width, and 500 μm in tip-to-tip spacing.

### cTnI Differential Aptalyzer fabrication

The Differential Aptalyzers were fabricated over a two-day process. Screen-printed electrodes comprising two central gold working electrodes (WEs), a surrounding silver reference electrode (RE), and an auxiliary gold counter electrode (CE) were used. The electrodes were cleaned with 2-isopropanol and deionized (DI) water, followed by electrochemical cleaning by cyclic voltammetry (CV) in 0.1 M sulfuric acid over a potential window of 0 to 1.5 V at a scan rate of 0.1 V/s. Each electrode underwent five CV cycles, with up to a maximum of 10 cycles performed, if needed, to match the currents of the paired working electrodes on a single chip. Cleaned chips exhibiting a reduction peak current < 35 μA or a current mismatch >5 μA between paired electrodes were excluded. Chips were thoroughly rinsed with DI water. For quality control, electrochemical characterization was performed in a redox buffer containing 1× PBS, 50 mM potassium chloride, 2 mM potassium ferrocyanide, and 2 mM potassium ferricyanide. Electrochemical impedance spectroscopy (EIS) was performed over a frequency range of 20 kHz to 0.5 Hz, and two CV scans between a potential range of −0.2 and 0.5 V at a scan rate of 0.1 V/s were performed.

Concurrently, free carboxyl groups of anti-cTnI and anti-BSA antibodies (0.086 μM each) were activated using EDC/NHS chemistry (40 mM, 20 mM, respectively) in MES buffer for 60 minutes. Cysteamine (0.6 μM) was subsequently added to the antibody solutions and incubated for 30 minutes. Antibody solutions were dialyzed for 60 minutes using 3F5 kDa molecular weight cutoff tubing to remove excess reagents. Dialyzed antibodies were immobilized onto freshly cleaned electrodes, with one WE modified with anti-cTnI and the second with anti-BSA on each chip. The chips were incubated overnight at 4 °C. The following day, electrodes were washed with 1× PBS and subjected to subsequent quality assurance scans to verify uniform antibody immobilization.

Thereafter, 3 μM thiol-terminated lactate aptamer was reduced with 200 μM TCEP for 2 hours at room temperature and subsequently deposited onto the chips. The aptamer immobilization was carried out for 2 hours at 4 °C, succeeded by an additional 2 hours of incubation at room temperature. The chips were then washed and dried prior to HMN patch attachment.

The crosslinked HMN patches were attached to the electrodes through a layer of 66.6 mg/mL MeHA solution. Briefly, 10 µL of the 66.6 mg/mL MeHA solution was drop-cast onto the four corners of the electrodes. Then, the HMN patches were gently placed on the electrodes to cover the counter, reference, and both working electrodes. Once the adhesive was completely dry, the Differential Aptalyzer was ready for subsequent testing.

### Swelling experiment

Porcine ear skin was cut into 1.5 cm × 1.5 cm squares and incubated in 1× PBS solution overnight to maintain hydration. The trimmed HMN patches were weighed (W□) and labeled prior to testing. Subsequently, the HMNs were applied and fixed onto the porcine skin for 2, 4, 8, 16, 32, and 64 minutes. After removal, the swollen HMNs were weighed again (W□). The swelling ratio was calculated using the following equation:

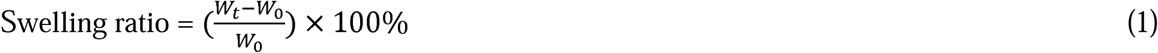

### *In vitro* evaluation of the differential chips (Differential Aptalyzer without the microneedles)

To evaluate the assay’s performance for cTnI detection, chips were exposed to a range of cTnI concentrations (0, 0.01, 0.04, 0.08, 0.16, 0.32, and 0.64 ng/mL) prepared in artificial ISF containing 2 mM or 4 mM lactate. Artificial ISF consisted of 2.5 mM CaCl_2_, 10 mM HEPES, 3.5 mM KCl, 0.7 mM MgSO_4_, 123 mM NaCl, 1.5 mM NaH_2_PO_4_, 7.4 mM sucrose, and 5.5 mM glucose. Target solutions (5 µL) were incubated on chip for 30 minutes prior to the measurement. Electrochemical sensing of the Differential Aptalyzer was performed using SWV in target solution, over the potential window of 0 to −0.5 V at 25 Hz (signal-off) and 100 Hz (signal-on). Measurements were acquired using PalmSens 4 potentiostat coupled to a MUX8-R2 multiplexer. Signal quantification was performed using the modified kinetic differential measurement (mKDM) method, calculated as the difference between the peak current measured at 100 Hz (signal-on frequency) and the peak current measured at 25 Hz (signal-off frequency), normalized by the peak current at 25 Hz.

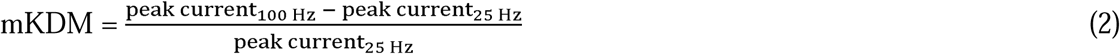

The acquired response was subsequently normalized with respect to the built-in lactate control at the baseline electrode and is expressed by equation (3) as follows:

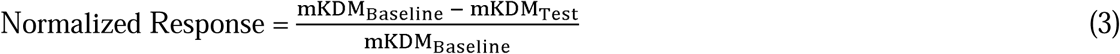

### *In vitro* validation of the cTnI Differential Aptalyzer

The cTnI Differential Aptalyzer was evaluated against varying cTnI concentrations (0, 0.01, 0.04, 0.08, 0.16, 0.32, and 0.64 ng/mL) prepared in 2 mM and 4 mM lactate, using the same protocols described above. Each cTnI Differential Aptalyzer was incubated with 100 μL of target solution for 30 minutes. Solution composition, instrumentation, SWV parameters, and signal quantification method were the same as above.

After obtaining Normalized Response curves, the limit of detection was determined at 2 mM and 4 mM lactate using the limit of blank (LOB) method^15^. The LOB was computed as mKDM_blank_ + 1.96 x σ_blank_, where σ_blank_ corresponds to the standard deviation of blank, defined as 2 mM or 4 mM lactate in absence of cTnI and 1.96 corresponds to the z-score for a 95% confidence interval. ^The LOD was calculated as LOB + 1.96 x σ^_lowest concentration_ ^where σ^_lowest concentration_ represents the standard deviation of lowest non-zero cTnI concentration dissolved in 2 mM or 4 mM lactate. Calibration curves were fitted using a non-linear one-phase association model. The limit of detection concentration was determined from the fitted model, mathematically defined as

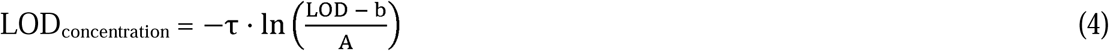

where τ in the inverse of the rate constant, b is the y-intercept of the fitted model, and A corresponds to the difference between plateau value and the y-intercept. The limit of quantification was calculated as LOB + 10 x σ_lowest concentration_ and corresponding limit of quantification concentration was determined using the same procedure as applied to the limit of detection concentration^51^.

### Mechanical test

The mechanical properties of the cTnI Differential Aptalyzer were tested using an Instron 5548 micro tester equipped with a 500 N compression cell. The cTnI Differential Aptalyzer patch was placed on the lower plate of the tester with the HMN tips facing upward. Before the test, the upper plate was positioned 1.5 mm above the needle tips. During the experiment, the upper plate descended at a constant rate of 0.5 mm/min, applying a vertical compression force to the HMN tips until it reached 70 N. The applied compression force and corresponding displacement of the HMN tips were recorded and plotted to determine the mechanical strength of the cTnI Differential Aptalyzer.

### Rhodamine B diffusion experiment

To visualize the diffusion process of molecules through the MeHA network, a Rhodamine B diffusion experiment was performed. MeHA HMNs were applied to a 1.4% agarose gel containing 7 µg/mL Rhodamine B for varying durations (10 s, 30 s, 1 minute, and 5 minutes). After each time point, the HMNs were removed, and the glass slides were immediately imaged using a fluorescence microscope (Nikon, Ti2), equipped with a TRITC filter set (excitation: 540–550 nm, emission: 565–620 nm). The fluorescence intensity within the microneedles was captured using an exposure time of 0.1 ms.

### Ferrocene-tagged protein diffusion

First, 400 µL of 150 µg/mL BSA was reacted with 2.5 µL of 10 mg/mL ferrocene-NHS dissolved in dimethylformamide (DMF). The reaction mixture was incubated at 4 °C for 2 hours, and the resulting ferrocene-labeled BSA (Ferrocene-BSA) was purified using an Amicon® Ultra centrifugal filter unit (30 kDa molecular weight cutoff). HMNs were then integrated with non-biofunctionalized dual-electrode chips as described above. The integrated HMNs were applied onto a 1.4% agarose gel containing 10 µg/mL Ferrocene-BSA. SWV measurements were performed every 5 minutes, starting 20 minutes after electrode application, over a potential range of 0 to 0.6 V at a frequency of 300 Hz and an amplitude of 100 mV, followed by three consecutive CV scans from 0 to 0.6 V at a scan rate of 0.5 V/s. Measurements were recorded for up to 45 minutes after HMN application.

### *In vitro* tracking of fluctuating cTnI levels

Differential chips functionalized with antibodies and aptamers were prepared and connected to the measurement setup. The chips were kept in a dark, humid environment, immersed in small beakers containing sufficient electrolyte to prevent drying. Transitions from artificial ISF to 4 mM lactate and subsequently from 4 mM lactate to 4 mM lactate + 0.08 ng/mL cTnI were achieved by spiking the electrolyte solution. To reverse conditions, the beakers were swapped to achieve transitions from 4 mM lactate + 0.08 ng/mL cTnI to 4 mM lactate, and subsequently from 4 mM lactate back to artificial ISF. In each condition, SWV measurements were collected for a period of 40 minutes with scans conducted every 10 minutes. Prior to each spiking event or beaker replacement, electrodes were subjected to pulse-assisted sensor regeneration protocol at a predefined potential and duration.

### Establishment of a mouse model of coronary artery disease

To validate the cTnI Differential Aptalyzer, we used a mouse model of coronary artery disease based on SR-B1−/−LDLR−/− mice, which are susceptible to diet-induced coronary artery atherosclerosis^46^. Male SR-B1−/−LDLR−/− mice were crossed with female SR-B1+/− LDLR−/− mice to generate littermate SR-B1+/−LDLR−/− controls and SR-B1−/−LDLR−/− disease mice. SR-B1−/−LDLR−/− mice were derived from breeders originally provided by Dr. Monty Krieger at the Massachusetts Institute of Technology. All animal procedures adhered to Canadian Council on Animal Care guidelines and were approved by the McMaster University Animal Research Ethics Board. Mice were bred and housed in the David Braley Research Institute facility with unrestricted access to standard chow (Teklad Global 18% Protein Rodent Diet, Envigo) and water. At 12 weeks of age, male and female SR-B1+/−LDLR−/− and SR-B1−/−LDLR−/− mice were switched to a high-fat, high-cholesterol, cholate-containing diet (HFHC; Teklad, TD88051; Envigo) containing 15% fat (7.5% from cocoa butter), 1.25% cholesterol, and 0.5% sodium cholate^52^ for 20 days to induce accelerated coronary artery atherosclerosis and myocardial infarction prior to the experiment.

### *In vivo* evaluation of cTnI Differential Aptalyzer using a mouse model of coronary artery disease

Mice were anesthetized with isoflurane (1.0–1.5% in oxygen at an oxygen flow rate of 1.0 L/min). Anesthesia was maintained for the duration of the procedures (approximately 3 hours from the start of fur removal to terminal blood collection and euthanasia). Throughout the procedure, mice were placed on a covered heating pad to maintain body temperature. Respiratory rate and heart rate were monitored regularly. Fur was removed from the backs of mice using Peanut clippers (Wahl Clipper Corporation) followed by a 10 min application of Nair depilatory cream. The skin was cleaned with a ChemWipe before two cTnI Differential Aptalyzers were applied to the skin using gentle thumb pressure and fixed in place using Tegaderm tape. After a 1-hour incubation, the ISF cTnI level was monitored for 60 minutes with a 10-minute interval using the cTnI Differential Aptalyzer. The results were reported as the average of the last five time-points. Following ISF measurement, blood samples were collected into a heparinized tube from the facial vein. Plasma was obtained by centrifugation of heparinized blood at 1200 × *g* for 10 minutes at 4 °C and subsequently aliquoted and stored at −80 °C. Plasma cTnI levels were quantified using the Ultra-Sensitive Mouse Cardiac Troponin I ELISA kit from Life Diagnostics, Inc. (Cat. No. CTNI-1-US). At the end of the experiment, the mice were weighed, euthanized under anesthesia by cervical dislocation, and hearts were removed, cryoprotected in 30% sucrose for 1–2 hours, embedded in O.C.T. compound (23-730-571; Fisher Scientific), snap frozen above liquid nitrogen, and stored at −80 °C for histological analysis.

For histological assessment of coronary artery atherosclerosis and myocardial damage, 10 µm transverse cryosections were obtained sequentially at 300 µm intervals from the mid heart to the aortic annulus and mounted on Superfrost Plus microscope slides (Fisherbrand, Fisher Scientific). Sections were either fixed in Bouin’s solution (HT10132, Sigma Aldrich) and stained with Masson’s Trichrome (HT15 1KT, Sigma Aldrich), or fixed in 37% formaldehyde and stained with Oil Red O (OD0395, BioBasic) and Mayer’s hematoxylin (1202C, Newcomer Supply) as described previously^46^. Bright-field images were captured using an Olympus BX41 microscope with a DP72 camera (Olympus Canada). For each heart, myocardial damage was evaluated in trichrome stained images of a section approximately 900 µm inferior to the aortic annulus. The percentage of myocardial damage was quantified in a blinded manner using the ratio of the area of myocardial damage relative to the total myocardial area of the cross section. Coronary artery atherosclerosis was evaluated blindly in 7 ORO-stained sections spaced 300 µm apart spanning 1800 µm below the aortic annulus. In each section, coronary artery cross-sections were categorized as non-occluded (0%), <50%, >50%, or fully (100%) occluded by lipid rich (ORO positive) atherosclerotic plaques. These scores were combined across all 7 sections, and the proportion of coronary artery cross-sections falling into each occlusion category was calculated per mouse.

### *In vivo* tracking of fluctuating cTnI with the Differential Aptalyzer

Human cTnI (T9924 Sigma Aldrich) or saline solution was injected into C57BL/6J wild-type mice (000664, Jackson Laboratory) to establish a cTnI fluctuation model. At the beginning of the experiment, mice were weighed and anesthetized, fur on the back was removed, and two cTnI Differential Aptalyzers were fitted as described. After one hour, single-frequency EIS was performed at 31.6 Hz, and the impedance magnitude (|Z|) was used for quality control. Differential Aptalyzers with external impedance (|Z| measured between the working and counter electrodes at 31.6 Hz) > 10 kΩ or internal impedance (|Z| measured between the two working electrodes at 31.6 Hz) > 2 kΩ were considered to have poor electrical contact and/or insufficient microneedle swelling and were excluded from subsequent analyses. Differential Aptalyzers that passed quality control were used to measure the baseline ISF cTnI level (prior to injection of human cTnI). Blood was collected (facial vein) into a heparinized tube prior to intravenous injection of human cTnI (50 μL in sterile saline) at doses of either 0 (saline alone), 5 or 25 μg human cTnI/kg body weight. ISF cTnI levels were monitored at 5, 15, 30, 45, 60, 75, and 120 minutes post-injection. Blood (facial vein) was collected into heparinized tubes at 5, 15, 60 and 120 minutes post human cTnI/saline injection. Plasma was prepared from heparinized blood as described above and plasma cTnI concentrations were measured using the Ultra-Sensitive Mouse Cardiac Troponin I ELISA kit from Life Diagnostics, Inc. (Cat. No. CTNI-1-US) using human cTnI as the standard, with values confirmed using the Human Cardiac Troponin I ELISA Kit from Abcam (Cat. No. ab200016). At the end of the experiment, the mice were euthanized under anesthesia by cervical dislocation.

## Acknowledgements

This work was supported by Ontario Early Research Award (MP), Johnson & Johnson WISTEM2d Award (MP), Canada Research Chair (MP & LS), Natural Sciences and Engineering Research Council of Canada (LS) and the Canadian Institutes of Health Research (PJT-178225 and PJT-195988 to BT). We also acknowledge the Translating Cardiovascular Remote Monitoring and Diagnostic Technologies for Health Equity (CaRDM Eq), a CREATE training program funded by Natural Sciences and Engineering Research Council of Canada.We thank Dr. Harmanjit Kaur and Dr. Bal Ram Adhikari for their fruitful discussion. Schematic figures were created using BioRender.com.

## Author contributions

H.Z., F.S., M.P. and L.S., B.L.T conceived the study. H.Z. and F.S. designed and performed the *in vitro* experiments. A.S.Q., B.L.T., M.P. and L.S. designed the *in vivo* experiments. H.Z., F.S. and A.S.Q. performed the *in vivo* experiments. All authors contributed to data analysis, figure creation and manuscript writing.

