## Supplementary Information for "Beyond Metabolites: A Wearable Differential Biointerface Integrating Antibody and Aptamer Probes for the Real-Time Tracking of Proteins In Vivo"

**for**

**Index**

**Figure S1.** Fabrication and characterization of Differential Aptalyzer

**Figure S2.** Normalized Response of the Differential Aptalyzer

**Figure S3.** Electrochemical signals recorded at signal off and on frequency in artificial ISF spiked with varying cTnI concentrations

**Figure S4.** 1H NMR spectra of methacrylated hyaluronic acid (MeHA)

**Figure S5.** Schematic illustration of HMN fabrication for ISF collection

**Figure S6.** Diffusion kinetics of ferrocene-tagged BSA through the Differential Aptalyzer

**Figure S7.** Pulse optimization of the cTnI Differential Aptalyzer

**Figure S8.** Increased coronary artery atherosclerosis in high-fat, high-cholesterol diet containing cholate (HFCC)–fed SR-B1-/-LDLR-/- mice

**Figure S9.** Magnified image of the dorsal skin of a mouse showing the insertion traces generated by the cTnI Differential Aptalyzer

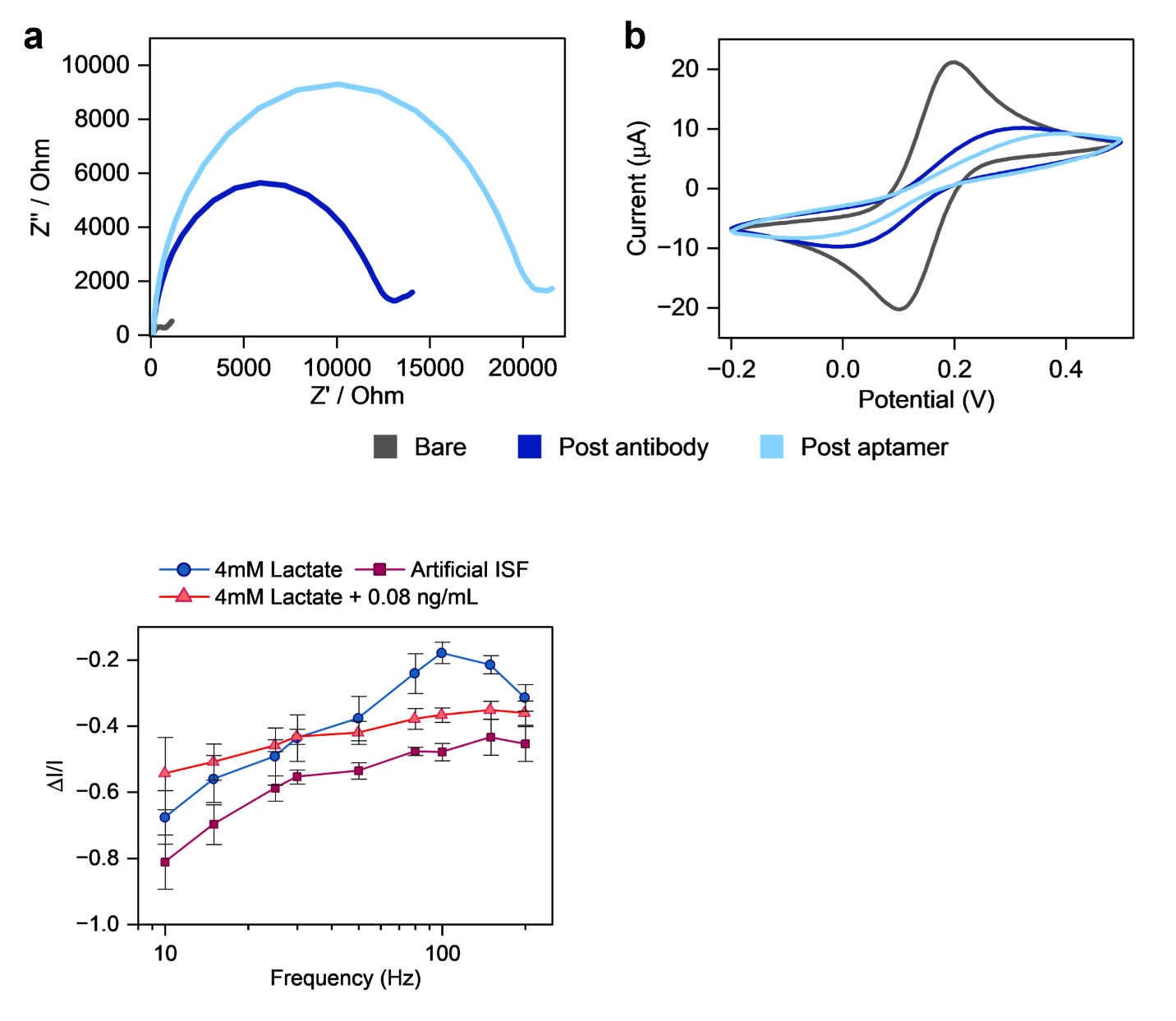

**Figure S1.** Fabrication and characterization of Differential Aptalyzer. **(a)** Electrochemical impedance spectroscopy (EIS) measurements were performed at frequency range from 0.5 Hz to 20000 Hz. **(b)** Cyclic voltammetry (CV) scans were performed at scan rate of 0.1 V/s and potential window of -0.2 to 0.5 V. All measurements are performed in 2 mM [Fe(CN)_6_]^3-/4-^ in 50 mM KCl and 1× Phosphate buffer Saline (1×PBS).

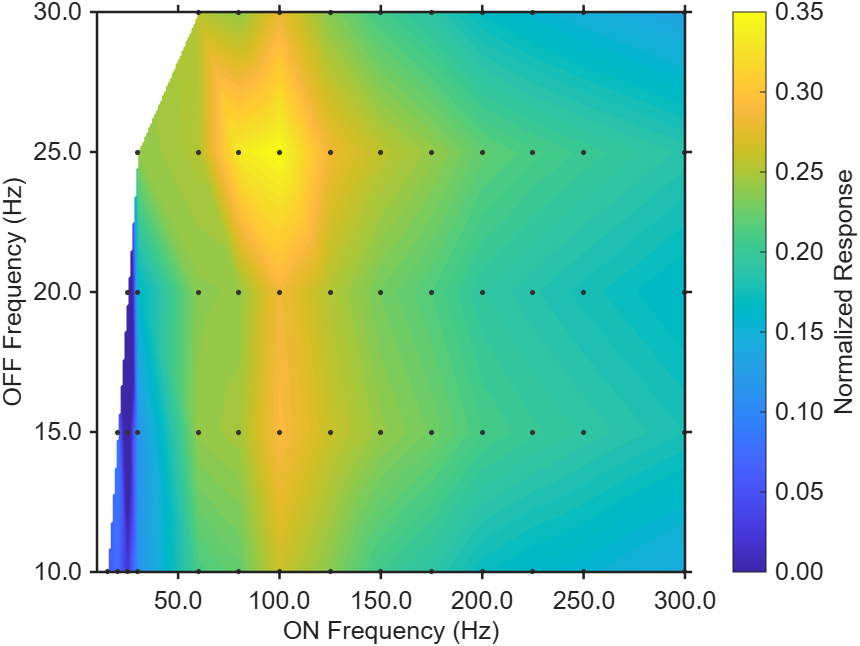

**Figure S2.** Normalized Response of the Differential Aptalyzer at various on and off frequencies at a cTnI concentration of 0.08 ng/mL and a lactate concentration of 4 mM. Measurements were conducted against Ag/AgCl reference electrode and Au counter electrode over a potential window of 0 to -0.5 V with 25 mV amplitude. The Black points indicate the specific frequency combinations that were tested.

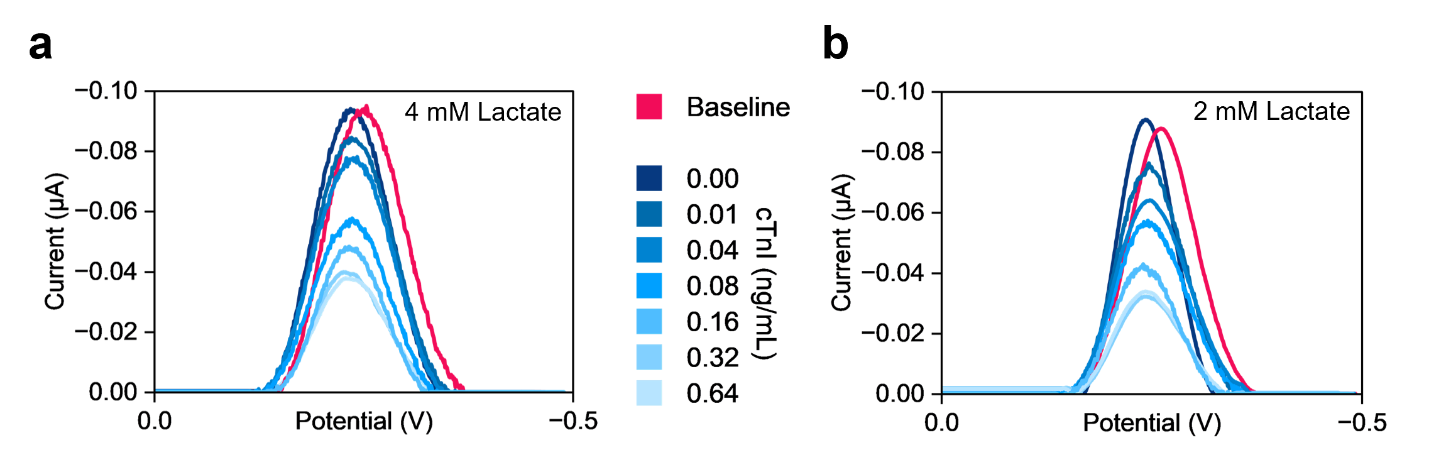

**Figure S3.** Electrochemical signals were recorded at 25 Hz off-frequency in artificial ISF spiked with varying cTnI concentrations, measured in **(a)** 4 mM and (**b)** 2 mM lactate. SWV were performed over a potential range of 0 to -0.5 V and an amplitude of 25 mV. The pink curve corresponds to the 25 Hz signal-off response obtained from the anti-BSA, baseline electrode, whereas the blue curves represent SWV responses acquired from anti-cTnI functionalized test electrodes. All SWVs were obtained from different chips, each exposed to varying cTnI in 2 mM or 4 mM lactate.

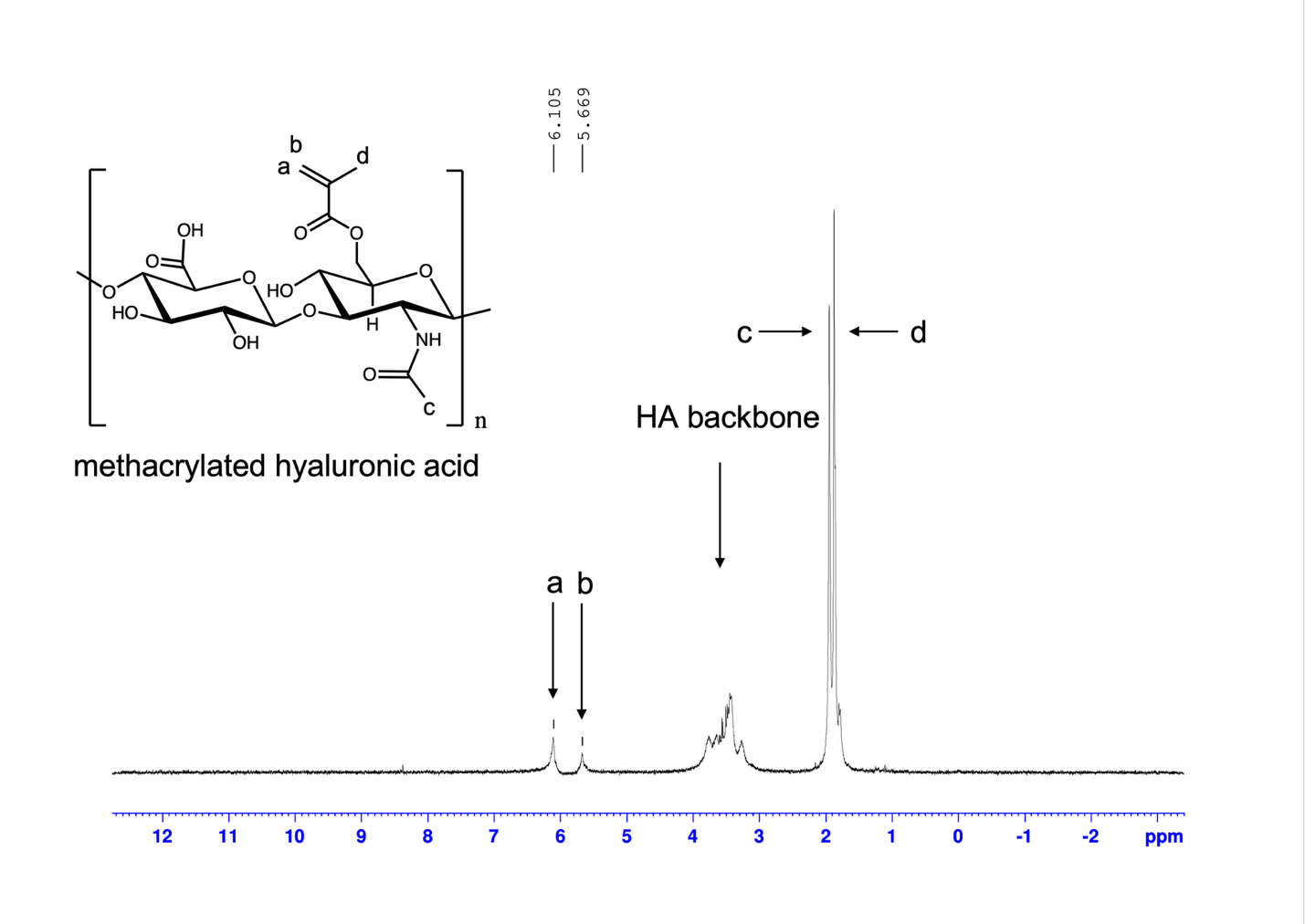

**Figure S4.** 1H NMR spectra of methacrylated hyaluronic acid (MeHA). MeHA was characterized with 300 MHz 1H NMR with a 10 ms-timescale to determine the degree of methacrylate modification. The appearance of the methacrylate vinyl resonances at δ ≈ 6.10 ppm (peak a) and 5.67 ppm (peak b) verifies successful functionalization.

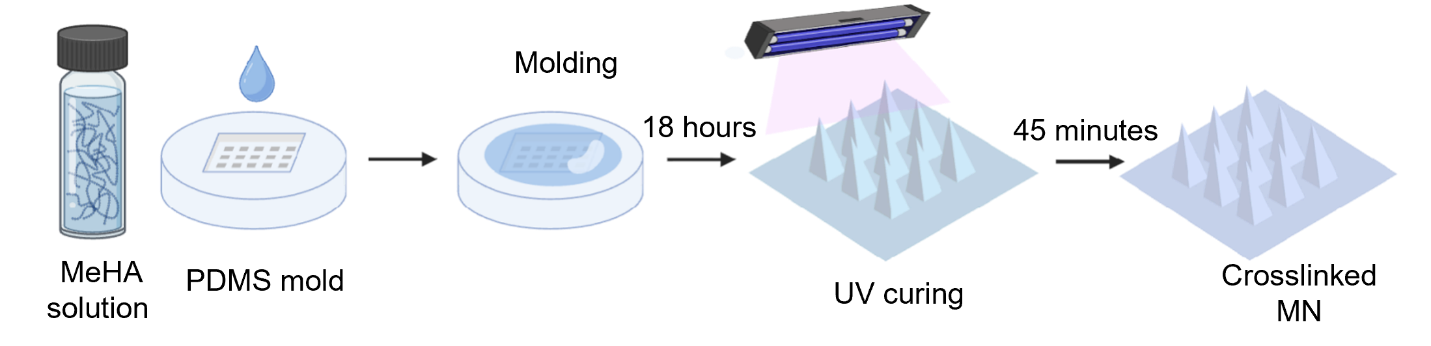

**Figure S5.** Schematic illustration of hydrogel microneedle (HMN) fabrication for interstitial fluid. A methacrylated hyaluronic acid (MeHA) precursor solution is cast onto a PDMS microneedle mold and drawn into the cavities by degassing. After overnight drying, the MeHA microneedle array is UV-crosslinked (~45 minutes) to yield a mechanically robust, crosslinked HMN patch ready for ISF collection.

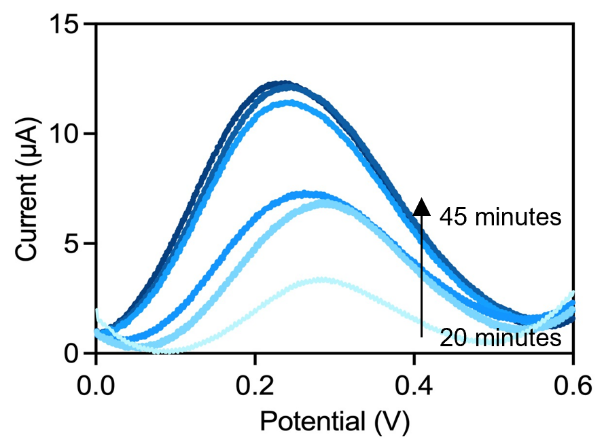

**Figure S6.** Diffusion kinetics of ferrocene-tagged bovine serum albumin (BSA) through the cTnI Differential Aptalyzer were evaluated by applying a 1.4% agarose hydrogel containing 10 µg/mL of ferrocene-tagged BSA to the hydrogel microneedle (HMN) side of the device and recording SWV at 20, 25, 30, 35, 40 and 45 minutes post-application. The SWV response reports the electrochemical signal of ferrocene-tagged BSA as it diffuses through the HMN matrix and reaches the working electrode surface. Measurements were performed using an on-chip Ag/AgCl reference electrode and an on-chip Au counter electrode over a potential window of 0 to 0.6 V at a frequency of 300 Hz and an amplitude of 100 mV.

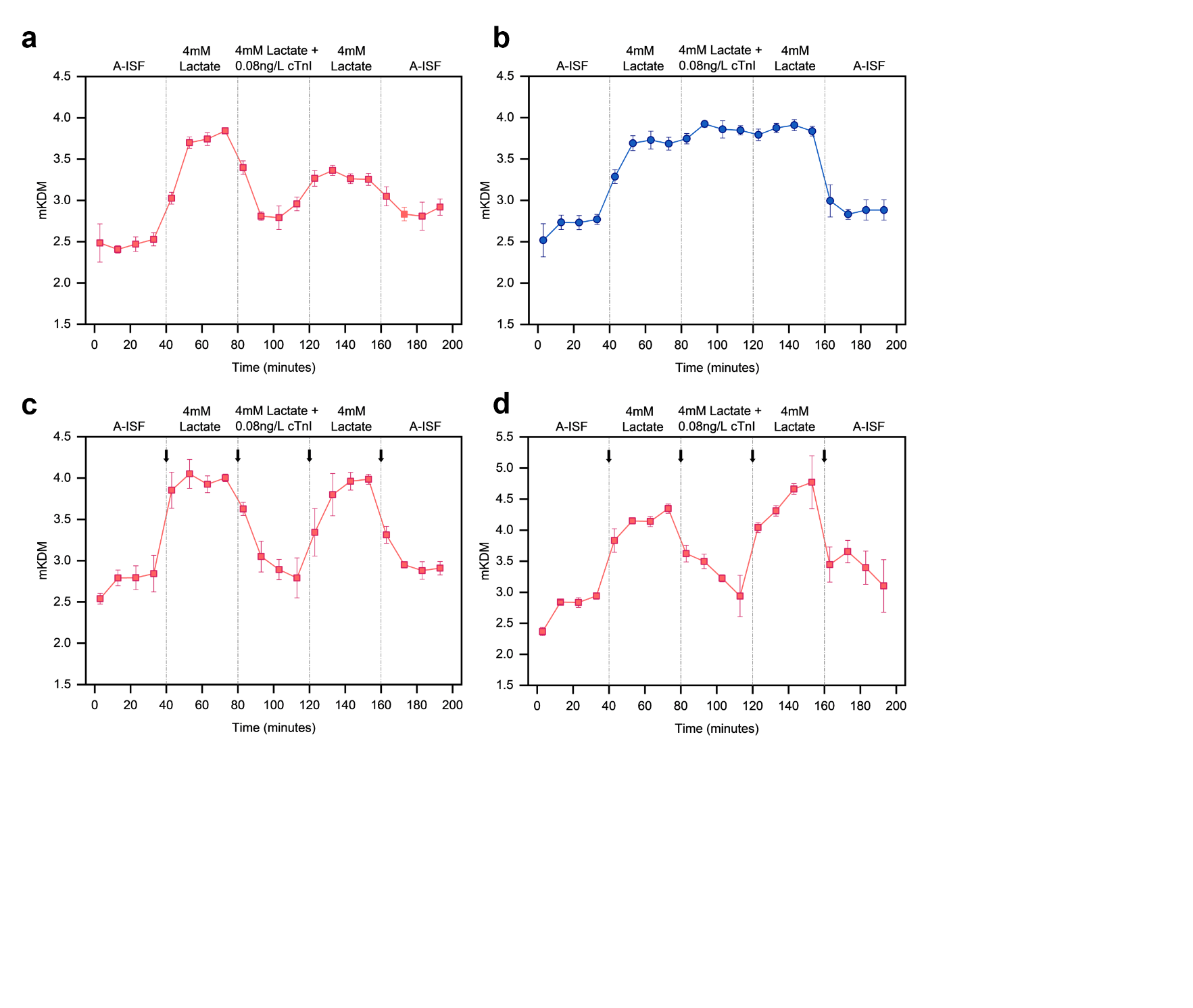

**Figure S7.** Optimization of the pulse-assisted sensor regeneration protocol for the cTnI Differential Aptalyzer. **(a)** The kinetic response of the test electrode in the absence of an applied pulse. **(b)** The kinetic response of the baseline electrode in the absence of an applied pulse. **(c)** The kinetic profile obtained following application of a -0.3 V pulse for 15 s to the test electrode. **(d)** The kinetic profile obtained following the application of a -0.3 V pulse for 15 s succeeded by a 0.3 V pulse for 15 s to the test electrode. The pulse was initiated at the timepoints indicated by black arrows.

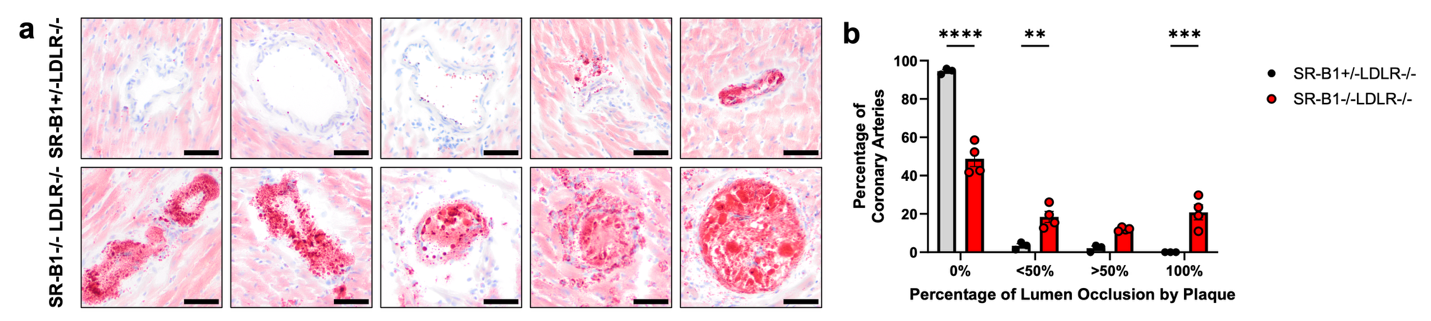

**Figure S8.** Increased coronary artery atherosclerosis in high-fat, high-cholesterol diet containing cholate (HFCC)–fed SR-B1−/−LDLR−/− mice. Serial cross-sections of hearts from the mid-ventricle through to the base of the aortic annulus were stained with Oil Red O and hematoxylin, and coronary arteries were scored as either 0% (nonatherosclerotic) or containing atherosclerotic plaques that occluded <50%, >50%, or 100% of their lumen. **(a)** Representative images of coronary arteries of increasing levels of occlusion from left to right from SR-B1+/−LDLR−/− (top) or SR-B1−/−LDLR−/− (bottom) hearts (scale bar = 50 µm). **(b)** Quantification of coronary arteries in each category across 7 heart cross-sections (n = 3, 4)^1^.

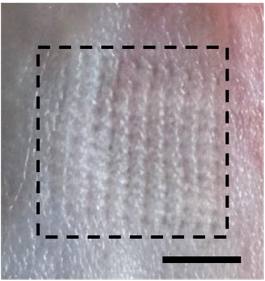

**Figure S9.** Magnified image of the dorsal skin of a mouse showing the insertion traces generated by cTnI Differential Aptalyzer, scale bar = 3.5 mm.
